# Neural correlates of perceptual switching while listening to bistable auditory streaming stimuli

**DOI:** 10.1101/669424

**Authors:** NC Higgins, DF Little, BD Yerkes, KM Nave, A Kuruvilla-Mathew, M Elhilali, JS Snyder

## Abstract

Understanding the neural underpinning of conscious perception remains one of the primary challenges of cognitive neuroscience. Theories based mostly on studies of the visual system differ according to whether the neural activity giving rise to conscious perception occurs in modality-specific sensory cortex or in associative areas, such as the frontal and parietal cortices. Here, we search for modality-specific conscious processing in the auditory cortex using a bistable stream segregation paradigm that presents a constant stimulus without the confounding influence of physical changes to sound properties. ABA_ triplets (i.e., alternating low, A, and high, B, tones, and _ gap) with a 700 ms silent response period after every third triplet were presented repeatedly, and human participants reported nearly equivalent proportions of 1- and 2-stream percepts. The pattern of behavioral responses was consistent with previous studies of visual and auditory bistable perception. The intermittent response paradigm has the benefit of evoking spontaneous perceptual switches that can be attributed to a well-defined stimulus event, enabling precise identification of the timing of perception-related neural events with event-related potentials (ERPs). Significantly more negative ERPs were observed for 2-streams compared to 1-stream, and for switches compared to non-switches during the sustained potential (500-1000 ms post-stimulus onset). Further analyses revealed that the negativity associated with switching was independent of switch direction, suggesting that spontaneous changes in perception have a unique neural signature separate from the observation that 2-streams has more negative ERPs than 1-stream. Source analysis of the sustained potential showed activity associated with these differences originating in anterior superior temporal gyrus, indicating involvement of the ventral auditory pathway that is important for processing auditory objects.

**Significance Statement:** When presented with ambiguous stimuli, the auditory system takes the available information and attempts to construct a useful percept. When multiple percepts are possible from the same stimuli, however, perception fluctuates back and forth between alternating percepts in a bistable manner. Here, we examine spontaneous switches in perception using a bistable auditory streaming paradigm with a novel intermittent stimulus paradigm, and measure sustained electrical activity in anterior portions of auditory cortex using event-related potentials. Analyses revealed enhanced sustained cortical activity when perceiving 2-streams compared to 1-stream, and when a switch occurred regardless of switch direction. These results indicate that neural responses in auditory cortex reflect both the content of perception and neural dynamics related to switches in perception.

## 1. Introduction

The moment-to-moment conscious states we all experience represent an enormous variety of experiences, due to our capacity to process many different types of stimuli while also incorporating internal and external contextual factors into our perceptual representations. According to the global workspace theory (Baars, 1988; Changeux and Dehaene, 2008; Dehaene and Changeux, 2011), individual sensory pathways process stimulus features unconsciously, until they arrive in frontal and parietal cortical areas that enable the widespread sharing of information about different features within and across modalities. In contrast, the information integration theory is more agnostic about where exactly in the cortex consciousness is generated, simply specifying that it can occur in any area that can generate different states in which the contents of awareness are integrated (Tononi et al., 2016). Still other theories say that basic forms of sensory awareness are generated in sensory cortex pathways, such as the ventral visual stream in the inferior occipital and temporal lobe (DiCarlo et al., 2012; Hochstein and Ahissar, 2002; Milner and Goodale, 2008; Pitts et al., 2012; Tong et al., 2006), without need for processing in associative areas such as the frontal and parietal cortex. Thus, there is still considerable debate about the neural basis of consciousness, including where in the brain it is generated. Moreover, almost all of these theories have been generated on the basis of visual studies, making it vitally important to also study auditory conscious processing to test the generality of these theories (Dykstra et al., 2017; Snyder et al., 2015).

Bistable stimuli provide an ideal means for experimentally manipulating consciousness because they induce mutually exclusive percepts that switch back and forth despite unchanging physical stimulus parameters. At the neural level, the standard model for bistable perception proposes that at any given time the current percept is destabilized over time due to adaptation, eventually reaching a threshold whereby the second percept becomes active and suppresses the first. Thus, competitive inhibition of both percepts results in a subjective experience of multiple percepts switching back and forth over time. In studies of binocular rivalry for example, two dissimilar images are presented simultaneously to each eye resulting in perception of one image or the other, spontaneously switching over time. Recordings of action potentials from individual neurons implicate the ventral visual pathway as the most likely locus for representation of the active percept (Leopold and Logothetis, 1996), although functional imaging studies in humans implicate earlier visual areas (Tong and Engel, 2001) and frontal and parietal networks (Lumer et al., 1998; Tong et al., 1998).

In both visual and auditory systems, there is ample evidence for diverging dorsal (“where”) and ventral (“what”) processing pathways (Arnott et al., 2004; Goodale and Milner, 1992; Lomber and Malhotra, 2008; Rauschecker and Tian, 2000). The ventral pathway, therefore, is a logical candidate for object identification, and in the case of complex scenes, segregation of separate objects and resolution of perceptual ambiguities. In the auditory literature, however, there is no evidence for involvement of the ventral pathway in bistable perception comparable to observations in the visual domain. Human imaging studies investigating bistable auditory stimuli have implicated primary and secondary auditory cortex in and around Heschl’s gyrus, as well as parietal cortical regions (Billig et al., 2018; Cusack, 2005; Gutschalk et al., 2008; Kondo et al., 2018). Moreover, the majority of these studies have focused on differences between the contents of perception or mechanisms of a switch in perception but not both (Kondo and Kashino, 2009; Sanders et al., 2018; Snyder et al., 2006). Finally, in both visual and auditory studies, the most direct evidence connecting neural adaptation and inhibition to perception comes from a study using magnetic resonance spectroscopy. Kondo et al., (2018) demonstrated a link between GABA/glutamate ratios in primary sensory cortices and percept duration during spontaneous fluctuations in perception, whereas prefrontal and parietal regions were linked to volitional control of perception. More specifically, the higher the GABA-to-glutamate ratio in frontal and parietal areas, the longer a percept was maintained, providing valuable insight into the neural dynamics between the active and alternate percept. However, questions remain about how the contents of perception are modulated, and how a perceptual switch is initiated relative to the global network responsible for conscious perception.

To answer these questions, we devised an experiment that uses an established bistable auditory stream segregation paradigm (Bregman, 1990; Van Noorden, 1975), but with intermittently presented stimuli (Kornmeier and Bach, 2004; Pitts et al., 2008). This paradigm presents triplets of ABA_ tones where A corresponds to a low tone, B to a high tone, and the blank, _, to the absence of a tone. When presented repetitively, these triplets can be perceived as either a single “galloping” auditory stream, or two separate streams (i.e., two metronomes). Typically, participants hold down one button (1-stream) or a second button (2-streams) to continuously indicate their perception. In this experiment, however, every third triplet is followed by a brief pause during which the participant presses the button to indicate their perception. The benefits of this approach are two-fold. First, it tightens the temporal link between components of the EEG and what a participant determines to be a 1- or 2-stream percept, potentially allowing for the separation of components related to the contents of perception and those related to the switch in perception. Secondly, it also changes the morphology of the event-related potentials (ERPs). In particular, the introduction of 700 ms of silence provides a well-defined baseline period, enabling clearer identification of the negative sustained potential (500-1000 ms), an auditory ERP that arises from the ventral auditory pathway that is linked to auditory object perception (Scherg et al., 1989). The sustained potential is therefore a component of the ERP expected to reveal effects of adaptation of the dominant percept according to standard theories of bistable perception (Brascamp et al., 2018; Rankin et al., 2015; Tong et al., 2006).

## 2. Materials and Methods

### 2.1. Participants

Thirty normal-hearing adults (11 male) with average age of 22.3 years (18-36 years) were recruited from the community in and around the University of Nevada, Las Vegas. All techniques and procedures were approved by the University of Nevada, Las Vegas Institutional Review Board. Experimental data, protocols, and analytical routines will be made available at https://osf.io/b4qrh/?view_only=81a1f5038e304822978d6d147ae70b3d, and upon direct request to the corresponding author. Prior to the experiment all participants provided informed consent followed by a standard hearing screening to ensure that audiometric thresholds did not exceed 25 dB hearing level at 0.25, 0.5, 1, 2, 4, and 8 kHz. An additional 23 individuals participated in the experiment were excluded due to a scarcity of trials in which a switch in perception was reported. Fourteen of these (out of N_total_ = 53) reported fewer than 20 total switches in perception throughout the experiment, overwhelmingly reporting 1-stream perception for the entire experiment. The remaining nine participants had noisy EEG data to a degree that less than 20 switch trials remained following the automatic epoch rejection that is described below. Therefore, the elevated number of excluded participants might be attributed to increased difficulty perceiving the 2-stream percept in this paradigm. In this case, our data would not provide insight for a subset of listeners who require different stimulus parameters to perceive 2-streams.

### 2.2. Intermittent Response Paradigm

A variation of the classic ABA_ auditory stream segregation paradigm was used. Participants were presented with repeating triplets of A and B tones, a stimulus that elicits alternating percepts of a single “galloping” auditory stream, or two separate “metronome” streams. Each 700 ms triplet consisted of A (400 Hz) and B (565.5 Hz) tones (6 semi-tone separation) presented in an ABA_ pattern with 175 ms separation between tone onsets, and a silent interval substituted for the 2^nd^ B tone (Fig. 1). Tones were 73 ms in duration. Three triplets presented in sequence followed by a 700 ms silent period designated for responding, defined each trial (2.8 s total). Prior to the experiment, participants were familiarized with the task and practiced conveying their perceptual response with a button press (button 1 for 1-stream, or button 2 for 2-streams; Cedrus response pad) during the 700 ms period following the 3 ABA_ triplets. The entire experiment was divided into 8 blocks of 75 trials presented in each block. Short breaks were provided to participants in between blocks.

**Figure 1.**
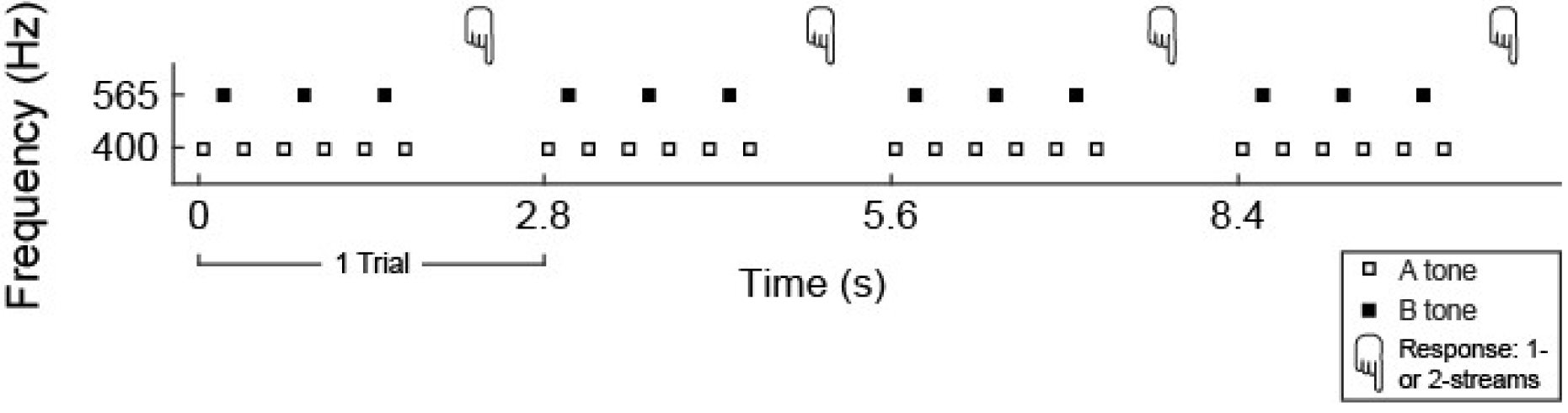
Stimulus presentation and intermittent response paradigm. Schematic depicting 4 trials of the intermittent bistable auditory stimulus. Each trial consisted of three triplets composed of pure tones presented in a low-high-low (ABA_) sequence. Participants indicated perception of 1- or 2-streams with a single button press during a 700 ms silent period at the end of each trial. Seventy-five consecutive trials made up an experimental block.

### 2.3. Stimulus Presentation

Auditory stimuli were presented to listeners via insert earphones (E-A-RTONE 3A Insert Earphones) at 65 dB SPL while sitting in a sound attenuation chamber. Prior to the experiment participants were instructed to keep their eyes focused on a white fixation cross on a gray background presented in the center of a computer screen and to report their perception with a button press. Participants were instructed to allow their perception to fluctuate without trying to hear the pattern one way or the other. All experimental stimuli were presented and responses recorded using routines written in the Julia programming language.

### 2.4. EEG data collection and analysis

During the task, EEG data were recorded using the BIOSEMI ActiveTwo system (512 Hz A/D rate) from 72 electrodes, including 8 external face electrodes used for artifact and eye-correction. EEG data were processed using EEGLAB (Delorme and Makeig, 2004) and custom Matlab routines. Individual participant data were referenced to the average of the two mastoid channels, bandpass filtered (0.01 – 30 Hz), and subjected to *infomax* independent component analysis (ICA) decomposition using the -extended and -runica options (Jung et al., 2000). The results were used to manually select and remove components related to ocular artifacts. Continuous data were then epoched for each 2.8 s trial and automatic epoch rejection (pop_autorej) was used to remove epochs that exceeded a threshold of 120 μV. Participants had to meet an inclusion criterion of at least 20 epochs retained that corresponded to a perceptual switch. Specifically, a participant must have indicated a switch in perception via button press in at least 20 trials, and at least 20 of those epoched trials must have survived automatic epoch rejection. Each epoch was then defined by percept and/or switch condition: 1-stream, 2-streams, switch, no-switch, switch from 1- to 2-streams, and switch from 2- to 1-stream. Trials designated as 1- or 2-streams did not include switch trials.

### 2.5. Statistical Analysis

Statistical comparison of waveforms was conducted using a non-parametric cluster-based analysis adapted from (Maris and Oostenveld, 2007). The first step in this process generated a test-statistic for each waveform-comparison by conducting a paired t-test across subjects comparing two conditions for each channel at each time point of the waveform (0-2100 ms; duration of sound presentation). Contiguous clusters of time points that exceeded the specified threshold of α=0.025 were identified and t-values within each cluster were summed together to create cluster-level statistics. The largest of the cluster-level statistics (summed t-values) for tested pair served as the test-statistic for comparison with a null-distribution (next step).

The second step generates a permutation, or null distribution by resampling each waveform-comparison via random partitioning; a process that in essence scrambles the condition-labels and resamples the data into two equal-sized new, or permuted datasets. This process was repeated 1000 times for each channel and subject, effectively resulting in 1000 resampled waveforms nominally corresponding to each condition. The outcome of this process generates 1000, 30-subject permuted datasets. The contiguous cluster analysis described above was then performed on each population, resulting in a permuted distribution of 1000 summed t-values representing the largest contiguous region of significance for each permuted population.

In the final step, a p-value was calculated based on the number of instances the permuted distribution from step 2 exceeded the test-statistic from step 1. If the probability was less than 0.05 (50 out of 1000), the difference was considered significant.

### 2.6. Source Analysis

Separate grand average waveforms (averaged across participants) for each of five conditions (all trials, 1-stream, 2-streams, switch 1-stream to 2-streams, switch 2-streams to 1-stream) were imported into BESA (Brain Electrical Source Analysis, Gräfelfing, Germany) software for dipole source analysis. The grand average that included all trials was used to find a general solution that accounted for the scalp data during the sustained potential using two pairs of symmetric dipoles. That solution was then applied individually to each of the other conditions, source waveforms were extracted for each of those conditions, and were used to qualitatively reconstruct individual comparisons observed in the scalp data.

## 3. Results

### 3.1. Behavioral Response Patterns

Response patterns reflecting perception of 1- or 2-streams were collected and analyzed from 30 participants. In an effort to establish that the intermittent presentation strategy employed here resulted in a similar pattern of bistable perception as the conventional continuous presentation paradigm, a number of perceptual characteristics were examined. First, in accord with previous studies, participants typically reported an initial bias to perceive 1-stream (Bregman, 1978; Pressnitzer and Hupé, 2006) followed by convergence towards an equivalent chance of reporting 1- or 2-streams (Sanders et al., 2018), approximately 10 trials into the block (Fig. 2A; black line). A similar measurement of switch probability measured over time revealed a consistent rate of switching around 0.2 (a switch was observed on about 20% of trials) over the time course of the blocks (total switches per participant: 106.5_mean_ +/-76_std_). In combination with the roughly equal probability of 1- versus 2-stream perception (1-stream probability: 0.55_mean_+/-0.1_std_), these results support the hypothesis that despite the intermittent nature of the paradigm, participants experienced stable perception over time, with occasional switches. If this were not the case, and the intermittent design failed to allow consistent perceptual buildup, the switch rate would likely be much higher, reflecting more frequent switches between the 1- and 2-streams percept due to interruptions in the ABA_ sequences (Cusack et al., 2004; Haywood and Roberts, 2013, 2010).

**Figure 2.**
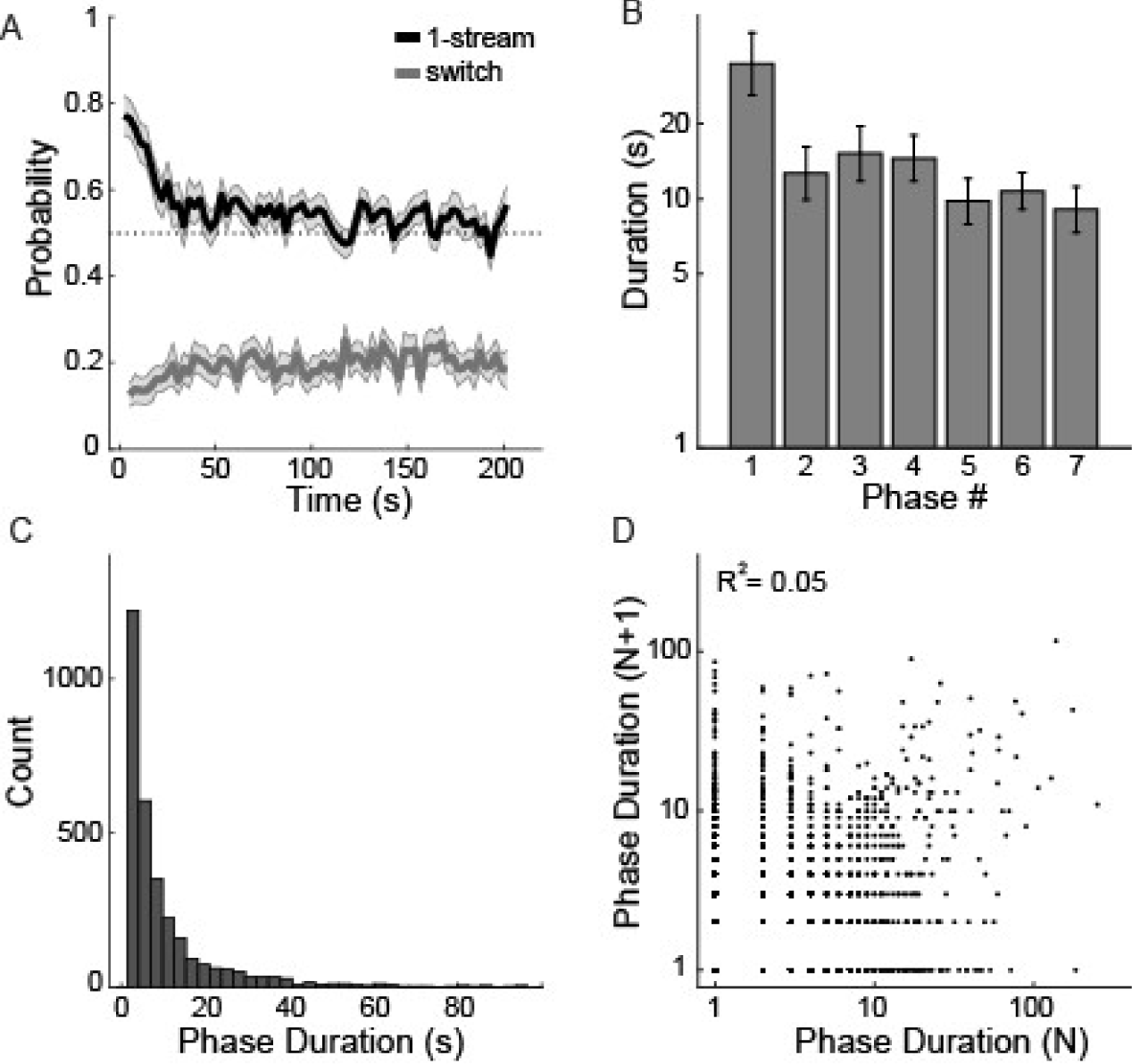
Behavioral characteristics of bistable perception. A) Probability of 1-stream perception (black line) and switch likelihood. Data represents 75 consecutive trials averaged across 8 blocks for each subject. B) The initial phase (length of time the same percept was reported) of each block exhibited longer duration than subsequent seven phases. Phase duration represents the number of consecutive 2.8 s trials the same percept was reported. C) The distribution of phase durations pooled across the participant-dataset approximates a logarithmic function, despite the discrete nature of the variable. D) Phase duration of a given percept (N) is minimally correlated (R2=0.05) with the phase duration of the next percept (N+1). Error bars in A and B indicate SEM across subjects (N=30).

Secondly, characteristics of the duration of a given percept – conventionally referred to as a “phase” – were also in agreement with prior research. Here, each phase was defined by the number of consecutive trials that the same percept was reported (each trial is 2.8 s). The initial phase of each block was significantly longer in duration than following percepts (Fig. 2B; repeated measures ANOVA: F_6,29_= 4.01, p<0.001, ŋ^2^_P_=0.12; post-hoc t-test phase 1 vs. phase 2: t_29_= 3.0, p<0.01, *d*=0.54), an observation believed to correspond to the build-up of segregation (Denham et al., 2013; Pressnitzer and Hupé, 2006). Due to the nature of the paradigm, phase measurements are necessarily a discrete variable with a minimum phase of 1 trial (or 2.8 s). Nevertheless, the distribution of phase durations approaches the shape of a logarithmic function (Fig. 2C), consistent with measures of bistable perception in both the auditory and visual domains (Farkas et al., 2018; Pressnitzer and Hupé, 2006). No significant difference was observed between the duration distributions for 1-stream versus 2-stream (Wilcoxon rank-sum test, Z=0.98, p=0.33). Lastly, the duration of a given phase (N) was not correlated with the duration of the following phase (N+1; Fig. 2D; R^2^ = 0.05).

### 3.2. ERPs: 1-stream vs. 2-streams

ERPs were grouped into categories corresponding to 1-stream or 2-streams, switch or no-switch trials. Switch trials were subsequently separated into those in which perception switched from 1- to 2-streams or switched from 2- to 1-stream. Presentation of long duration auditory stimuli evokes a sustained negative potential that appears at frontal electrodes (Picton et al., 1978a, 1978b), and is localized to an area anterior to the portion of auditory cortex that generates the N1 (Scherg et al., 1989). This sustained potential was observed in ERPs during perception of both 1- or 2-streams, but had larger amplitude in response to 2-streams compared to 1-stream (Fig. 3A; top row). Significant differences were observed at FC2 and F1, and overall enhanced negativity for 2- versus 1-stream at channels surrounding Cz (Fig. 3A; topography). This pattern of results is consistent with a number of EEG and MEG studies (Billig et al., 2018; Gutschalk et al., 2005; Snyder et al., 2015, 2009) using comparable ABA_ paradigms, with the prevailing explanation that a 2-stream percept represented by separate neural populations evokes greater activity at scalp electrodes. Note that significant differences were observed over a longer time range than indicated by the gray shading in Figure 3A, but those differences did not reach the conservative contiguity requirement of the analysis.

**Figure 3.**
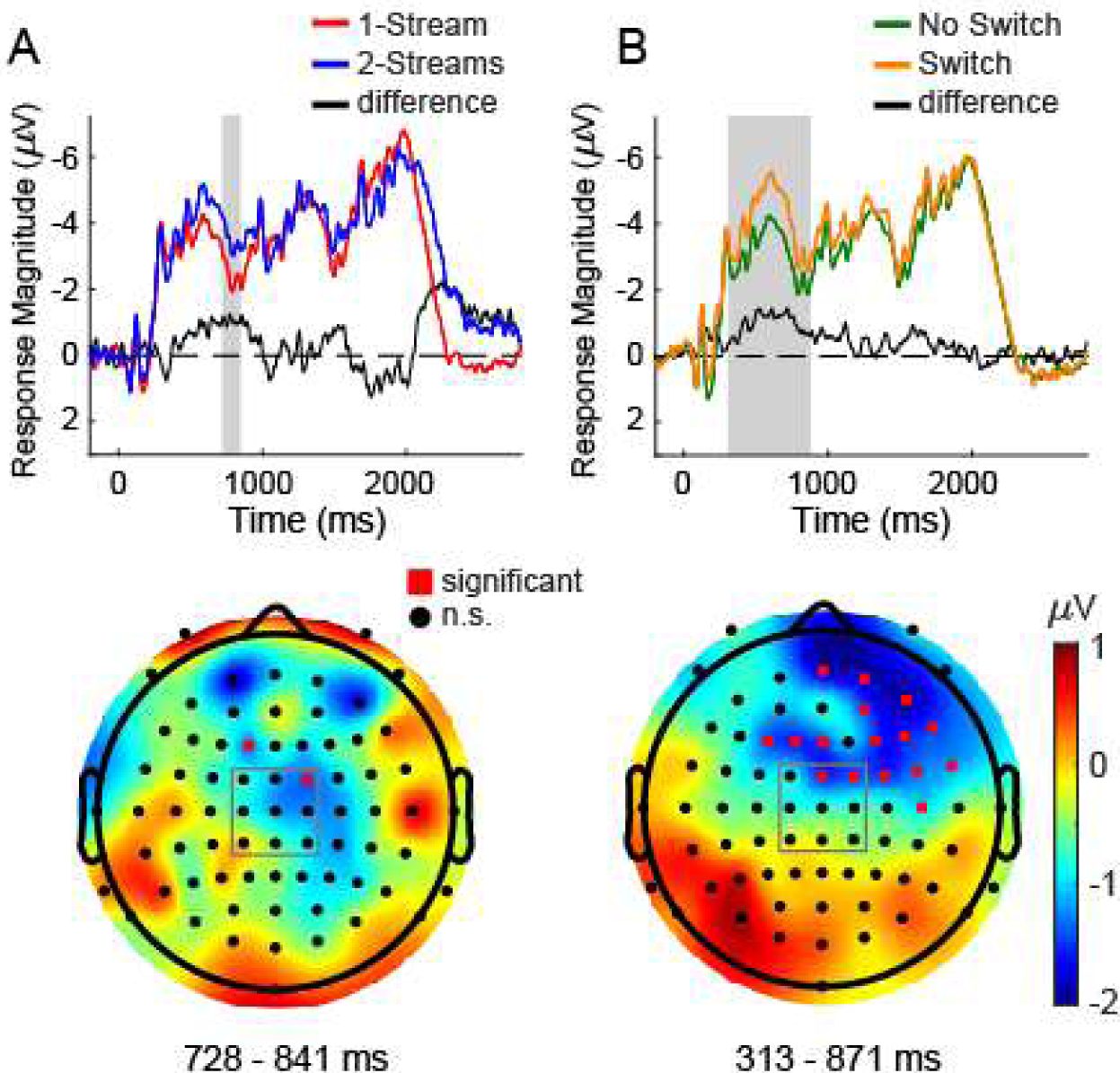
Enhanced negative responses observed during the sustained potential for comparisons of A) stable 2- versus 1-stream percept, B) switch versus no-switch in perception. Top row: average ERP waveforms for the 9 channels surrounding Cz (black square on topographic map) and the difference wave for each comparison (black line). Grey shading indicates time range of contiguous significance for individual channels. Bottom row: spatial organization of channels overlaid on topographic map of the difference between conditions. Red circles indicate location of channels with significant differences.

### 3.3. ERPs: switch vs. no-switch

The last trial of each phase by definition was a trial in which a switch in perception must have occurred. ERPs corresponding to switch trials were compared to no-switch trials using the cluster-based permutation described above. The results revealed significantly greater negative responses in the sustained potential across multiple channels for switch compared to no-switch trials (Fig. 3B; top). Differences were mainly located at right-frontal electrodes, as reflected in the difference topography (Fig. 3B; bottom). There are two potential reasons for this observed difference. The switch versus no-switch comparison does not distinguish between switching from 1- to 2-streams or switching from 2 to 1 stream, and could therefore reflect the fact that perceiving 2 streams results in larger activity, described above, regardless of the fact there was a switch. Alternatively, the observed differences could be due to switching independent of the percept. To address these possibilities two additional analyses were conducted. First, a comparison of switch type was made between switches from 1- to 2-streams and a switch from 2- to 1-stream (Fig. 4A). This revealed a temporal-spatial deviation, in which significant differences for three channels located on the top of the head were significantly different during an early part of the waveform (45-175 ms), whereas a large number of leftward channels displayed significant differences during a later part of the waveform, temporally similar to the comparisons shown in Figure 3. These observations suggest at least part of the switch versus no-switch difference is attributable to enhanced negativity associated with the perception of 2-streams versus 1-stream. Secondly, two additional comparisons were made in an effort to identify an effect of switch versus no-switch, while controlling for the already established effect of percept. Trials with a switch from 2-streams to 1-stream were compared to stable (no-switch) 1-stream percepts (Fig. 4B), and those with a switch from 1- to 2-streams were compared to stable (no-switch) 2-stream percepts (Fig. 4C). In both cases, channels with significantly more negative potentials were observed for switch trials during the sustained potential portion of the ERP at frontal electrodes.

**Figure 4.**
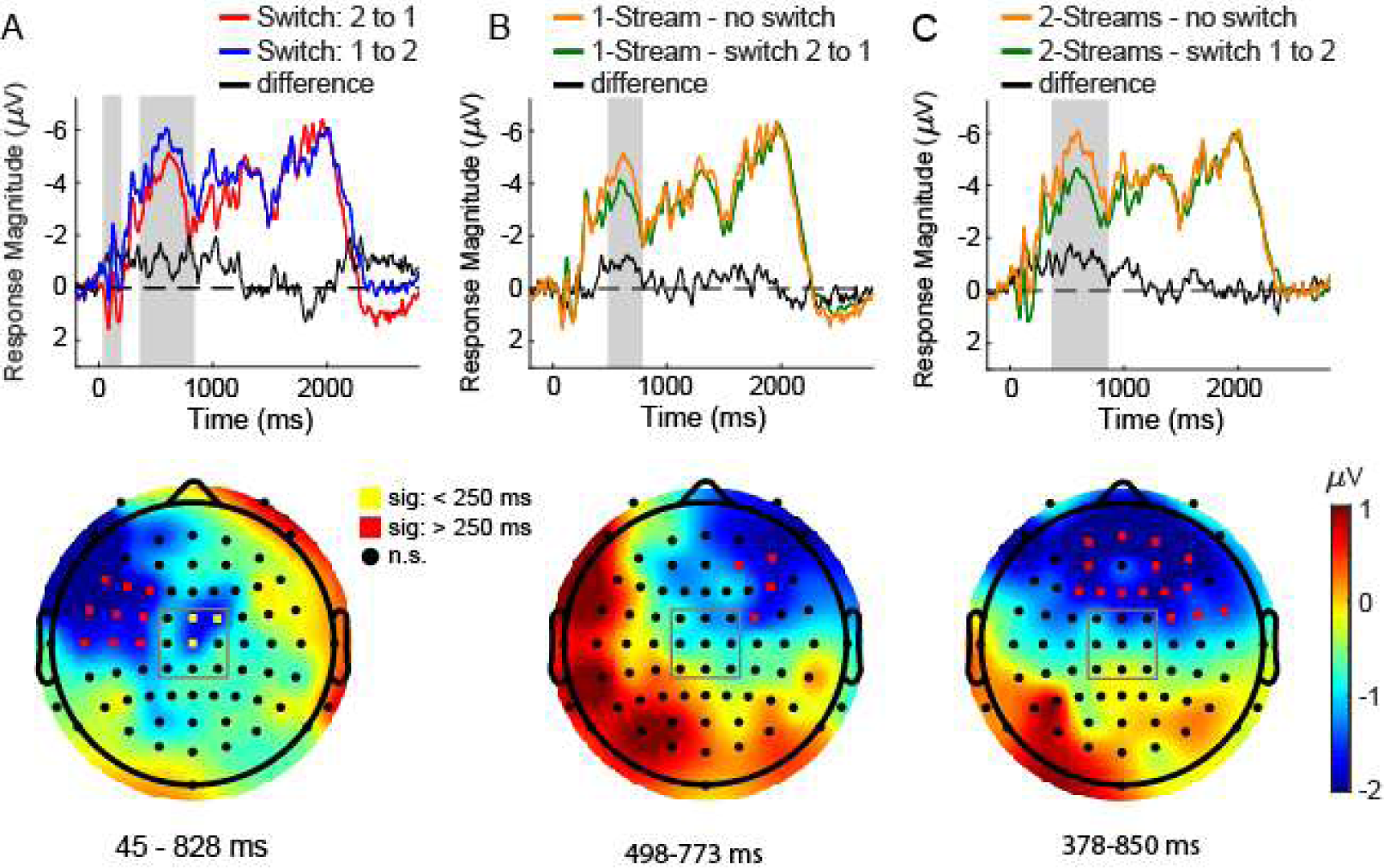
Enhanced negative responses observed during the sustained potential for comparisons of A) switch from 1- to 2-streams versus 2- to 1-stream percept, B) switch from 2- to 1-stream versus stable 1-stream percept, C) switch from 1- to 2-streams versus stable 2-stream percept. Top row: average ERP waveforms for the 9 channels surrounding Cz (black square on topographic map) and the difference wave for each comparison (black line). Grey shading indicates time range of contiguous significance for individual channels. Bottom row: spatial organization of channels overlaid on topographic map of the difference between conditions. Colored circles indicate location of channels with significant differences observed at earlier (yellow, 50 - 200 ms) and later (red, 200 - 800 ms) time ranges.

### 3.4. Source Analysis

Symmetric pairs of dipoles located bilaterally in auditory cortices and parietal lobes (Fig. 5A) accounted for 96.1% of the variance in the scalp data observed across all trials measured over a large portion of the epoch (100 – 2100 ms post-stimulus). The time range includes transient responses (N1, P2) as well as the later sustained potential, encompassing all three triplets of the trial. This solution was applied to each of the individual conditions retaining the original dipole orientations and over the same time range: 1-stream, 2-stream, switch 1 to 2, and switch 2 to 1 and in all cases explained a large proportion of the variance (Explained Variance > 0.87; Table 1) for each condition. Source waveforms qualitatively replicated the results presented in figures 3 and 4 (Fig. 5C; top row): 2-stream activity was greater than 1-stream, and switching conditions had more activity in the sustained potential than non-switching conditions (Fig. 5B). The dipoles in auditory cortex alone also accounted for a large portion of the overall sustained potential response variance (Explained Variance > 0.86; Fig. 5C Table 1), but source waveforms isolated from these dipoles alone poorly reflect the difference between 1- and 2-stream percepts observed in Figures 3A and 4A (Fig. 5C; middle row). Interestingly, differences between 1- and 2-stream percepts is best reflected in the parietal sources, specifically located in medial parietal cortex. The closest cortical areas to these sources are precuneus and posterior cingulate cortex, regions associated with Gestalt-type integration of features into coherent objects (Pflugshaupt et al., 2016), and the dorsal attention network (Raichle et al., 2001), respectively.

**Table 1.** Source analysis on waveform representing all trials yielded a 4 dipole and 2 dipole solution. This solution, when applied to individual conditions accounted for the indicated percentage of the variance. Dipole numbers correspond to labels in Figure 5.

| Condition | Variance Explained |  |
| --- | --- | --- |
|  | Dipoles 1-4 | Dipoles 1-2 |
| All Trials | 96.12 | 94.74 |
| 1-Stream | 91.04 | 89.08 |
| 2-Streams | 87.05 | 85.61 |
| Switch 1 to 2 | 91.44 | 88.62 |
| Switch 2 to 1 | 89.25 | 86.76 |

**Figure 5.**
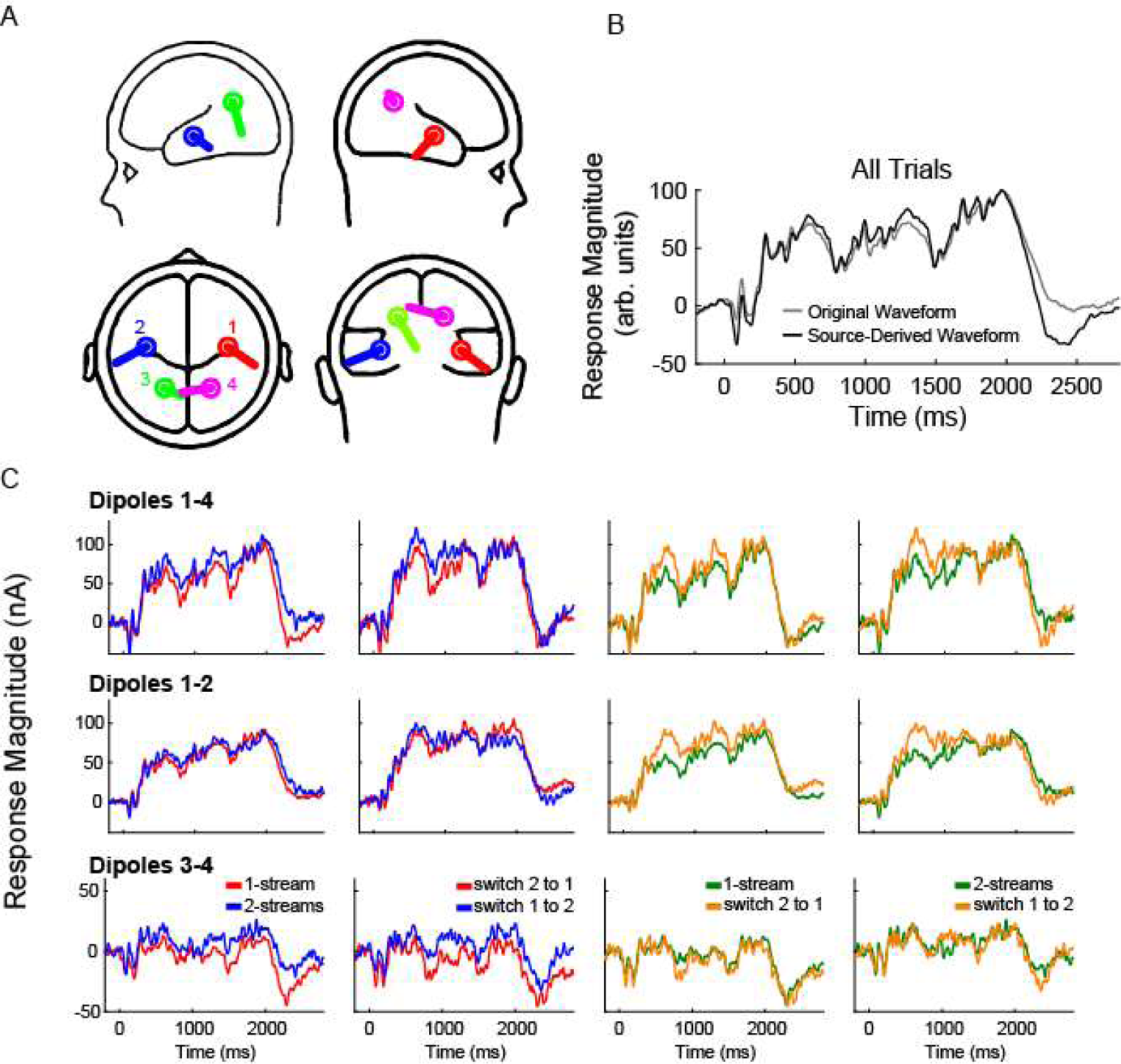
Source analysis of the sustained potential. A) Symmetric dipoles located in left (blue) and right (red) hemisphere anterior STG and left (green) and right (cyan) hemisphere of the parietal lobe. B) GFP (Global Field Power) of the original waveform including all trials of the experiment (gray) and the source-derived waveform (black) based on a time window from 100-2100 ms. GFP of source waveforms for individual conditions corresponding to dipoles 1-4 (top row), auditory cortex dipoles 1-2 (middle row), and parietal dipoles 3-4 (bottom row).

## 4. Discussion

To explore stable and dynamic aspects of conscious auditory perception, we performed an intermittent ABA_ auditory streaming experiment. Presenting the auditory stimuli in relatively discrete segments helped us identify modulations of the sustained potential during a switch in perception compared to stable periods, independent of switch direction. The sustained potential also reflected the contents of perception, namely whether participants were perceiving one vs. two streams, during the stable periods.

### 4.1. Behavioral Response Patterns

The ABA_ streaming stimulus has been used extensively for experiments on auditory scene analysis and is commonly conducted using one of two general approaches. The first is a continuous-presentation design in which participants constantly indicate perception via button press over the time-course of multiple minutes (Anstis and Saida, 1985; Carl and Gutschalk, 2013; Denham et al., 2018; Pressnitzer and Hupé, 2006). The second typically consists of a two-part sequence with an induction period followed by a test period in which a manipulation of the ABA_ stimulus along one or more dimensions (e.g., temporal, spectral, location) serves as a probe for perceptual continuity (Haywood and Roberts, 2010; Rogers and Bregman, 1993; Yerkes et al., 2019). The first approach accommodates for the observation of spontaneous switching of perception over an extended period of time, while the second provides a better-defined event associated with a perceptual switch. Despite the temporal discontinuity, the current findings follow established behavioral patterns characteristic of the continuous button-response paradigm: balanced time for each percept (Fig. 2A), an initial percept of 1-stream characterized by longer duration (Fig. 2A, 2B), a logarithmically shaped distribution of phase duration (Fig. 2C), and lack of correlation between sequential phase durations (Fig. 2D). As a result, the intermittent-response paradigm tested here incorporates benefits from each paradigm type, the observation of spontaneous switching behavior over time, and a well-defined stimulus event for linking perception to modulations of ERPs.

### 4.2. Sustained Potential

Most of the ERP differences observed between 1- versus 2-streams (Fig. 3A, 4A) and switch versus non-switch conditions (Fig. 3B, 4B, 4C) were over the portion of the waveform considered to be the auditory sustained potential. This brain response is characterized by negative voltage at frontal scalp locations in response to continuous auditory stimulation (Kohler and Wegener, 1955; Picton et al., 1978a, 1978b; Scherg et al., 1989). Unlike earlier responses to sound onsets and offsets that exhibit more transient positive and negative deflections in the 75-200 ms range, the sustained potential is unaffected by mixed presentations of click and tone-burst stimuli, and in the context of auditory stream segregation has been shown to be sensitive to attention and features of the ABA_ tones such as frequency separation (Snyder et al., 2006).

In prior work, source analysis of the underlying neural generators of the sustained potential revealed bilateral, vertically oriented dipoles in anterior superior temporal gyrus (STG) (Scherg et al., 1989). In agreement with this finding, the current study found optimized dipoles located bilaterally in anterior STG. Anterior STG is a region linked to the “what” part of the “what/where” dual-pathway model for sound pattern identification (Ahveninen et al., 2013; Bizley and Cohen, 2013; Rauschecker and Tian, 2000; Zündorf et al., 2016). Thus, our results support the hypothesis that the differences observed in the sustained potential relate to the neural signature of the streaming process, whereby stimulus features are integrated/segregated to determine perception of 1- or 2-auditory streams. Evidence for separate processes is observed in the ERPs, where differences between switch and no-switch trials appear earlier and extend over a longer time range compared to differences between representation of 1- and 2-stream perception (Fig. 3A). This timing difference is also apparent in the source analysis where dipoles 1-2 corresponding to auditory cortex showed greater separation for switch versus no-switch conditions earlier in the waveform (Fig. 5C; middle row), whereas dipoles 3-4 corresponding to parietal sources showed the most separation for 1- versus 2-streams later in the waveform (Fig. 5C; bottom row), consistent with fMRI studies (Cusack, 2005; Teki et al., 2011). In summary, the ERP differences presented here suggest separate processes underlie the neural signatures for the 1- versus 2-stream difference and the switch versus no-switch difference. Whether there are separate mechanisms that modulate a switch from 1- to 2-streams or 2- to 1-stream should be addressed by future experiments.

### 4.3. Implications for theoretical models of bistability

Separating target information from background noise is one of the primary tasks of a sensory system. In the face of ambiguous stimuli, the conventional dynamic model for bistable perception proposes that populations of neurons representing different states incorporate inhibition, adaptation, and noise to generate bistablity or multistablity (Brascamp et al., 2018; Rankin et al., 2017, 2015; Tong et al., 2006). Inhibition leads to competition between different states, and adaptation and noise allow for switches between the currently dominant state.

During stable periods of perception, such a model may show large, unadapted responses. As a steady percept is maintained, adaptation reduces responses until a threshold is crossed, at which time the non-dominant percept overtakes the dominant percept and a switch in perception is triggered. Interpretation of the results of this study within the framework of this model is fairly straightforward. The large potentials observed at the very beginning of a percept, while termed a “switch” in this experiment, also correspond to a fresh percept prior to the forthcoming effects of adaptation. As the percept proceeds, adaptation builds up until a switch occurs and another fresh percept (with a large ERP) emerges.

From a modeling perspective, such a sequence of events could be used to differentiate switches from non-switch periods on the basis of the amount of adaptation and inhibition present. The present finding that the signature for switches began earlier in the waveforms and in more sensory regions than the recognition of a 1-stream vs. 2-stream perception (Fig. 3A compared to 3B; Fig. 5C, bottom row compared to middle row) has implications for the locus of sources of adaptation, inhibition and noise that may drive bistability. Several existing models of bistability in auditory streaming are consistent with an early locus for bistability (Rankin et al., 2017, 2015), while others assume bistability occurs as part of the process of identifying the number of sources in an auditory stream (Barniv and Nelken, 2015; Mill et al., 2013); these latter models do not appear to be consistent with our data because they predict a similar locus for switching and recognition of 1-vs. 2-stream percepts. Note that none of the existing auditory models account for the effects of the 700 ms break present in the current study, but could easily be modified on the basis of computational studies that do consider the effects of gaps (Noest et al., 2007; Vattikuti et al., 2016). In summary, models that allow for early sensory sources of bistability (Noest et al., 2007; Vattikuti et al., 2016) or those that posit some form of top-down modulation of sensory competition (Brascamp et al., 2018; Kleinschmidt et al., 2012; Li et al., 2017) appear most consistent with our data.

### 4.4. Conclusion

In this study, we present data from an auditory streaming experiment using an intermittent stimulus paradigm that showed behavioral characteristics consistent with continuous bistable perception while maintaining control of the temporal dynamics important for recording ERPs. Consistent with previous studies, sustained auditory potentials associated with perception of 2-streams exhibited greater negative potentials than 1-stream. Unexpectedly, sustained potentials were significantly more negative when a perceptual switch occurred, regardless of the switch direction, leading to the conclusion that perceptual switches have a neural correlate unique from the overall representation of 1-stream or 2-streams. Importantly, the ability to tease apart the neural correlates associated with a) an internally derived event (a switch in perception) and b) an ongoing perceptual representation, can be attributed to the unique intermittent design employed in this experiment.

## Acknowledgements

Supported by Office of Naval Research: N00014-16-1-2879

